# Phage-Inducible Chromosomal Jumbo Islands (PICJIs) spreading antimicrobial resistance, host-adaptation factors and defense systems, a novel satellite family of nucleus-forming *Chimalliviridae*

**DOI:** 10.64898/2026.11.03.763896

**Authors:** Aurore Rossignol, Karine Praud, Mélodie Roxane Alexandre, Myriam Mikhayel, Isabelle Foubert, Pierre-Ivan Raynal, Laetitia Fabre, Sébastien Olivier Leclercq, François-Xavier Weill, Dolla Karam Sarkis, Benoît Doublet

## Abstract

Phage satellites are mobile genetic elements that depend on helper bacteriophages to promote their own dissemination. Here, we identified a novel family of large satellites, named Phage-Inducible Chromosomal Jumbo Islands (PICJIs), that are widespread in *Escherichia coli* and *Salmonella enterica* among diverse hosts, worldwide. These largest satellites to date (71.6 ± 15.8 kbp) are specifically transduced by the persistent jumbo bacteriophage *Goslarvirus* of the nucleus-forming *Chimalliviridae* family. Furthermore, PICJIs carried numerous accessory genes associated with clinically relevant antimicrobial resistances, host-adaptation, and several anti-phage defense systems.

## Main text

Phage satellites constitute a large family of integrative mobile genetic elements that depend on a helper bacteriophage for their own life cycle^1^. An increasing number of new types of satellites have been recently described, the most well-characterized families being P4-like, phage-inducible chromosomal islands (PICIs, i.e. SaPIs of *Staphylococcus aureus*), and the PICI-like elements (PLEs) of *Vibrio cholerae* (for review see ref.^1^)^2,3^. In the absence of an active helper phage, most satellites remain passively integrated within the bacterial chromosome. The interactions between satellites and helper phages extend from parasitism with some satellites hijacking the helper machinery and interfering with phage reproduction (selfish parasites of parasites) to a mutualistic relationship ultimately beneficial both for the helper phage and the bacterial host^1,4,5^. Whilst satellites are generally small in size (∼15 kbp), the transfer of larger genomic islands carrying defense systems by a ‘jumbo’ nucleus-forming phage (i.e., characterized by double-stranded DNA genomes exceeding 200 kbp) have been recently described in *Pseudomonas*^6,7^.

From previous studies^8,9^, we identified two unrelated *Escherichia coli* strains (B25FEP and M24FEP) isolated from healthy broilers in Lebanon (Supplementary Table 1) carrying a novel 72.1-kbp transferable integrative element, hereafter named EcCI^B25FEP^ (For *E. coli* Chromosomal Island of strain B25FEP), harbouring the extended-spectrum β-lactamase gene *bla*_CTX-M-55_ and five other antimicrobial resistance (AMR) genes (Fig. 1a). Mating experiments and genomic analyses of B25FEP, M24FEP and their related *E. coli* transductants revealed that EcCI^B25FEP^ integrates specifically at the 3’-end of *tRNA-Phe*, resulting in 19-bp direct repeats *attL* and *attR* flanking EcCI^B25FEP^ (Fig. 1a, Extended Data Fig. 1). In addition to numerous accessory genes described in more detail below, EcCI^B25FEP^ lacks recognizable genes encoding conjugative transfer machinery (e.g., mating pore formation, relaxase, and Type 4 coupling protein), but exhibits common features of phage satellites (Integrase-, excisionase-, and primase-encoding genes) such as PICIs, cf-PICIs (capsid forming PICIs), P4-like satellites, PLE, and PICMIs (minimalist PICIs) except its larger size and was not detected by SatelliteFinder^1–5,10–12^. Recently, numerous jumbo phages notably of the nucleus-forming family *Chimalliviridae* were categorized as persistent virulent phages able to persist in *E. coli* cultures despite streaking on plates when isolating strains and without strong host mortality^7,13,14^. We thus hypothesized that the field strains B25FEP and M24FEP initially harboured a helper jumbo phage able to transduce the large chromosomal island EcCI^B25FEP^ as described in *Pseudomonas*^6^. Three independent phage lysates were produced from the animal *E. coli* strain B25FEP and assessed in EcCI^B25FEP^ transduction assays (see Methods). All lysates enabled the EcCI^B25FEP^ transduction into the recipient *E. coli* DH5α with various efficiencies ranging from 6.0 x 10^3^ to 1.6 x 10^5^ EcCI^B25FEP^ transductant forming units (TFUs) per ml of phage lysate (Extended Data Fig. 2). The integration of EcCI^B25FEP^ at the 3’-end of *tRNA-Phe* was confirmed by PCR in all tested transductants. In parallel, lysis efficiency on *E. coli* DH5α of these lysates was found to be correlated with the transduction efficiency, suggesting that the putative helper phage was able to replicate in *E. coli* DH5α (Extended Data Fig. 2).

**Fig. 1.**
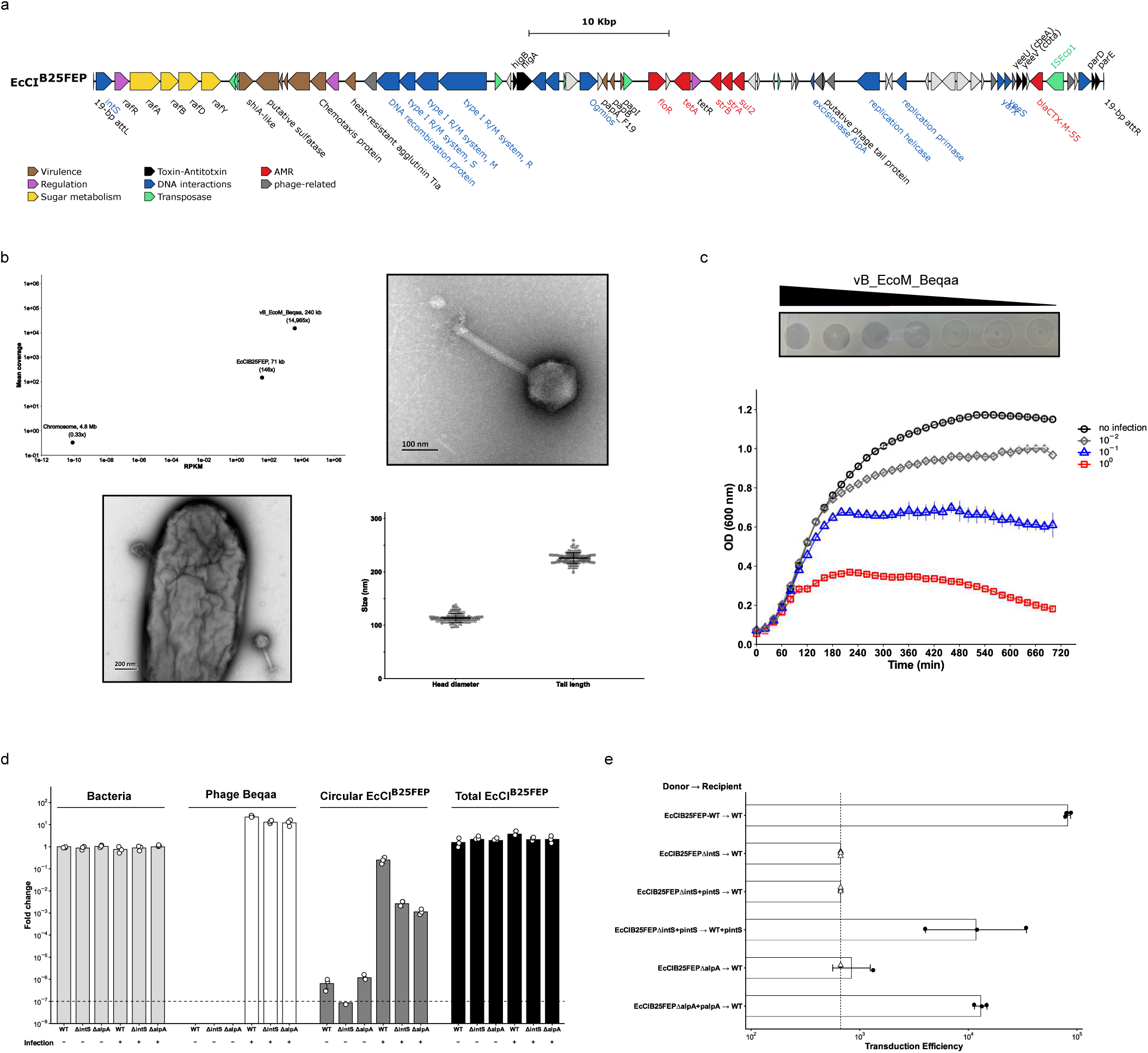
Transduction of the chromosomal island EcCI^B25FEP^ by the jumbo phage vB_EcoM_Beqaa. **a**, Genetic organization of the *tRNA-Phe*-integrated chromosomal island EcCI^B25FEP^. Open reading frames are color-coded according to protein functions in legend. Relevant gene names or functions are indicated on the map. The left and right 19-bp direct repeat sequences *attL* and *attR*, respectively, flanking EcCI^B25FEP^ are indicated by thick black bars. **b**, The upper-left panel shows the scatter plot of the relationship between RPKM (Reads Per Kilobase per Million mapped reads) and mean coverage of Illumina mapped reads for the phage vB_EcoM_Beqaa, the EcCI^B25FEP^, and the *E. coli* chromosome in whole phage DNA of B25FEP lysate. Mean coverages are indicated for each point. (see also Extended Data Fig. 4b for ONT mapping data). Transmission electron micrographs of a vB_EcoM_Beqaa phage particle from NaCl/PEG-purified lysate (upper-right image) and from *E. coli* B25FEP culture infected by vB_EcoM_Beqaa (bottom-left image). Samples were negatively stained with 1% uranyl acetate. Scale bars are included in each picture (see also Extended Data Fig. 5). The bottom-right panel illustrated morphological dimensions of vB_EcoM_Beqaa phage particles. Individual measurements are represented by grey dots. Horizontal bars indicate the mean and error bars denote standard deviation. A total of 100 phage particles were analysed among hundreds visualized. Raw uncropped images and source data are provided as Source Data file. **c**, vB_EcoM_Beqaa phage infection of *E. coli* DH5α. Upper panel, Plaque assays of 10-fold serial dilutions of phage Beqaa lysate on *E. coli* strain DH5α. Lower panel, Growth curves of *E. coli* DH5α cultures infected by 10-fold serial dilutions of phage vB_EcoM_Beqaa. Each curve represents the mean ± SD of five independent experiments. (see also Extended Data Fig. 6 for EcCI^B25FEP^-carrying *E. coli* DH5α, Source Data file). **d**, qPCR quantification of the circular form of EcCI^B25FEP^ wild-type and single-gene mutants (Δ*intS* and Δ*alpA*) with or without vB_EcoM_Beqaa infection. Fold change was calculated by comparing DNA samples using ΔΔCt to the bacterial chromosome of the non-infected wild-type arbitrary placed at 1. Source data are provided as Source Data files. The dashed line indicates the limit of detection for this assay. Bars represent the geometric mean ± geometric standard deviation (GSD) from three independent experiments (individual dots). **e**, Transduction efficiency of EcCI^B25FEP^. Phage lysates were produced on different donor strains carrying EcCIB^25FEP^ (wild-type, Δ*intS*, Δ*alpA*, and *in-trans* complementation) and used to infect recipient *E. coli* strains DH5α. Transductants were selected on ceftriaxone and transduction efficiency was expressed as TFU.ml^-1^ of phage lysate. Bars represent the geometric mean ± GSD from three independent experiments (individual dots). The dashed line indicates the limit of detection for this assay (667 TFU.ml^-1^). Triangles represent experiments arbitrary placed at the detection threshold for which no transductant was obtained.

To identify the helper phage, the whole DNA of B25FEP lysate was extracted after DNase treatment and sequenced using short- and long-read technologies. Hybrid assembly returned a 240,606-bp virulent phage sequence without identity with EcCI^B25FEP^ but belonging to the *Chimalliviridae* family and sharing 96.7% average nucleotide identity with the nucleus-forming jumbo *Goslarvirus Escherichia* phage vB_EcoM_Goslar which have been isolated from duck feces in Germany in 2016 (Extended Data Fig. 3)^15^. Since, the phage Goslar became a model among the *Chimalliviridae* family that forms a proteinaceaous phage nucleus and a cytoskeletal filament vortex during host infection^16,17^ (for recent reviews see ref.^7,18^). Accordingly, this novel jumbo *Goslarvirus Escherichia* phage was named vB_EcoM_Beqaa related to its geographical origin in Lebanon (hereafter referred to as Beqaa for simplicity). To date, Beqaa constitutes the second isolated *Goslarvirus* in addition to 37 *Goslarvirus* genomes identified in *Enterobacteriaceae* assemblies^13^. Moreover, *Goslarvirus* belonged to persistent virulent phages recently described to be able to persist in *E. coli* cultures without strong host mortality^13,14^, being consistent with our hypothesis that the phage Beqaa was present within *E. coli* cultures and concomitantly isolated from the gut microbiome of healthy broilers^8^. Mapping of sequencing reads obtained from DNA extracted from purified particles against the complete genomes of Beqaa and EcCI^B25FEP^ revealed the presence of complete EcCI^B25FEP^ in phage particles (38.38 RPKM), but with a mean coverage ∼100-fold lower than the genome of the helper phage Beqaa (3927 RPKM) that was also confirmed by qPCR (Fig. 1b, Extended Data Fig. 4). In addition, the few reads mapping the *E. coli* chromosome (8.23 10^-11^ RPKM) showed characteristics of highly fragmented sequences, suggesting that they may correspond to chromosomal DNA fragments randomly packaged before lysis or to genomic DNA incompletely digested by DNase from the supernatant. Our sequencing results strongly suggest that EcCI^B25FEP^ is preferentially packaged compared to the chromosome by the jumbo phage Beqaa in a way similar to phage satellites^1,4^. Moreover, EcCI^B25FEP^ has a genome one third smaller than the size of its helper phage Beqaa, a common rule among families of known phage satellites^1^. Transmission electron microscopy of B25FEP lysate showed that the helper phage has a typical myovirus morphology with a large icosahedral head and a long contractile tail composed of a tube and a sheath (Fig 1b, Extended Data Fig 5), consistent with the jumbo *Goslarvirus* morphology^19^. Beqaa particles have a constant capsid diameter and tail length of ∼ 114 ± 8 and ∼ 226 ± 10 nm, respectively (Fig. 1b, Extended Data Fig 5)^7,15,19^. The analysis of hundreds of particles did not reveal any smaller-sized capsids, suggesting that contrary to cf-PICIs, EcCI^B25FEP^ does not produce small capsids exclusively packaging its own genome^11^.

To further confirm these results, colonies of the original *E. coli* strains B25FEP and M24FEP were successively restreaked on plates until being fully depleted of the jumbo phage Beqaa to obtain pure bacterial strains carrying the integrated EcCI^B25FEP^ element^14^. The purified clones were negative for Beqaa by PCR and deficient for the EcCI^B25FEP^ transfer in transduction assays. In parallel, the jumbo phage Beqaa of the initial phage lysate from B25FEP was also purified and amplified on the sensitive *E. coli* strain DH5α to obtain pure Beqaa lysate without EcCI^B25FEP^-containing particles (see Methods and Supplementary Table 1). In spot assays, the phage Beqaa produced a complete clearing until the 10^-1^ dilution and partial lysis at higher dilutions but without apparent plaques (Fig. 1c, Extended Data Fig 6). We then challenged *E. coli* DH5α harbouring or not EcCI^B25FEP^ with the phage Beqaa at three different MOI (pure lysate to 10^-2^ dilution) over 12h infection assays. The jumbo phage Beqaa inhibited the growth of *E. coli* DH5α in a MOI dependent manner showing only slight growth inhibition at the 10^-2^ dilution in this condition (Fig. 1c). Similarly to the phage Goslar, Beqaa infection does not strongly affect the bacterial host population that is consistent for phages which persist within their hosts^14^. In addition, no significant difference of susceptibility to Beqaa was observed between *E. coli* DH5α harbouring EcCI^B25FEP^ and the control strain (Extended Data Fig. 6).

We then decided to investigate the direct induction of EcCI^B25FEP^ by the helper phage Beqaa. The integrase gene *intS* and the gene *alpA* encoding for a putative excisionase were deleted in EcCI^B25FEP^, and the mutants were compared to the wild-type element for excision and transduction. In absence of Beqaa infection, EcCI^B25FEP^ showed a very low level of excision (10^-6^) that is fully abolished by the Δ*intS* deletion, but remained still detectable for the Δ*alpA* mutant by qPCR (Fig. 1d). The Beqaa infection strongly induced excision of the wild-type EcCI^B25FEP^, in ca. one quarter of the cells in these assays. Moreover, Beqaa infection confirmed that both genes *intS* and *alpA* are essential to achieve a high level of EcCI^B25FEP^ excision induced by the helper phage (Fig. 1d). The *intS* and *alpA* deletions also led to transduction efficiencies below and very close to the detection limit, that are consistent with no detection and ∼ 2 log decrease of the amount of circular EcCI^B25FEP^ by qPCR in the corresponding phage lysates, respectively (Fig. 1e, Supplementary Table S4). Plasmidic complementation of *intS*, under the control of an inducible promoter, was necessary both in the donor and recipient strain to restore the transduction of EcCI^B25FEPΔ^*^int^* (Fig 1e). In contrast, *in trans* complementation of *alpA* was sufficient in the donor to induce EcCI^B25FEP^ excision and to restore in part the EcCI^B25FEPΔ^*^alpA^* transduction (Fig 1e). Together, these results confirmed that the excision of EcCI^B25FEP^ is induced by the infection of its helper phage Beqaa. Due to its large size and specific induction by a helper Jumbo phage, we classified EcCI^B25FEP^ among a novel family of satellites that we named PICJIs for Phage-Inducible Chromosomal Jumbo Islands including the recently described I55 element of *Pseudomonas* also induced by the jumbo phage JPTI55^6^.

Blast search in prokaryotic complete genomes revealed that the EcCI^B25FEP^ element was a representative of a diverse family of *tRNA-Phe* integrated elements in *E. coli* and *Salmonella enterica*. A total of 273 complete EcCI^tRNA-Phe^ elements were identified in diverse *E. coli* from all phylogroups and in few *S. enterica* strains, mainly of the serotype Indiana (belonging to ST17) (Supplementary Table 2). All these strains were isolated between 1984 and 2025 from various human, animal, food and environmental sources, worldwide (Supplementary Table 2). To further confirm the restricted host range of EcCI^B25FEP^, transduction assays were realized in different *Enterobacterales* genera and species (Supplementary Table 1). We only succeeded to transduce EcCI^B25FEP^ into the reference *S.* enterica serotype Indiana strain 573K (ST17) at low frequency (10^3^ TFU per ml of phage lysate) close to the detection limit in repeated attempts (Supplementary Table 1, Extended Data Fig. 1). However, phage lysates produced from Beqaa-infected *S.* Indiana tranductant 573K-TDT were negative for EcCI^B25FEP^ transduction. In addition, Beqaa plaque assays with *S.* Indiana strain 573K and its transductant 573K-TDT show a very slight lysis strongly suggesting a poor replication of the jumbo phage Beqaa into *Salmonella* (Extended Data Fig. 6). EcCI^B25FEP^ transduction into *S.* Indiana probably constitute an example of “silent” transfers across genera that are rarely demonstrated experimentally^1^.

Overall, these 274 EcCI^tRNA-Phe^ elements are highly diverse in size ranging from 13.7 to 118 kbp (avg. 71.6 ± 15.7 kbp) as well as in gene content. Sequence analysis using PPanGGOLiN tool identified 9 soft-core genes notably encoding the tyrosine integrase (*intS*), the excisionase gene (*alpA*), the DNA primase (*pri*) and the TA system (*parDE*) that are shared among 237 EcCI^tRNA-Phe^ elements. A phylogenetic tree was built to infer their evolutionary relationship based on the alignment of these 9 soft-core genes (Fig. 2a, Extended Data Fig. 7). The largest cluster (n=121) containing EcCI^B25FEP^ encompasses nearly all elements carrying AMR genes with a great majority highly-related to EcCI^B25FEP^. The two other large clusters (n=53 and 47) are more diverse genetically as well as in size and lacks AMR genes. Fifteen EcCI^tRNA-Phe^ elements representative of the diversity were fully aligned for detailed comparison (Fig. 2b). EcCI^tRNA-Phe^ elements appeared to be preferential hotspots for horizontally-acquired genes mediated by different IS elements. Beside AMR, the main accessory determinants are the raffinose metabolism operon *rafRABDY* carried by 247 out of 274 elements and the virulence-associated genes *shia* and *tia* found in 227 and 179 elements, respectively (Fig. 2a, Extended Data Fig. 7, Supplementary Table 2). The raffinose metabolism conferred by *rafRABDY* was confirmed by growth experiments. EcCI^B25FEP^-carrying *E. coli* DH5α were able to grow in minimal medium supplemented with raffinose as sole carbon source while no growth was observed for the recipient *E. coli* DH5α (Fig. 2c). Raffinose being a prebiotic, this metabolic advantage could play a role in host adaptation contributing to nutrient competition with other beneficial bacteria and consequently to a better colonization of the intestinal tract^2,4^.

**Fig. 2.**
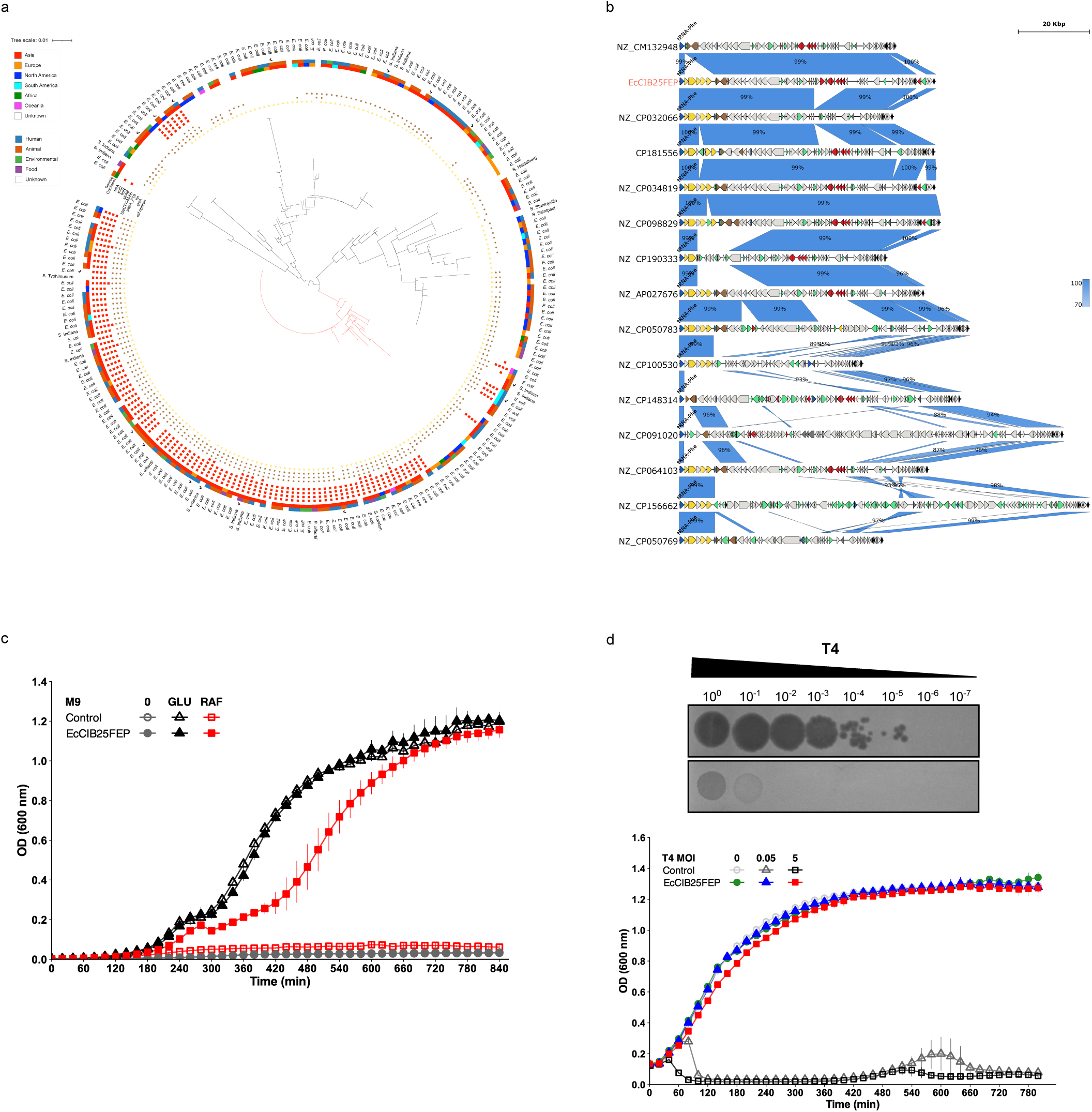
Diversity of EcCI^B25FEP^-related chromosomal islands and functional characterisation of EcCI^B25FEP^. **a**, Phylogenetic relationship of 237 EcCI^B25FEP^-related chromosomal islands integrated at *tRNA-Phe* (Supplementary Table 2). The tree is based on the alignment of soft-core genes (see methods) and rooted at mid-point. Branches of the largest cluster containing EcCI^B25FEP^ are coloured in red. Source, geographical origin, bacterial host species, presence/absence of the raffinose metabolism operon, main virulence-associated genes, AMR genes and size are shown from inside to outside of the tree. Genetic organisation of EcCI^B25FEP^-related elements marked with a tick are further compared in panel b. Additional core genes are also highlighted in Extended Data Fig. 7. **b**, Complete genetic maps and comparisons of 15 representatives of the EcCI^B25FEP^ family. Open reading frames are color-coded according to protein functions as in Fig. 1a. Sequence homologies larger than 500 bp are shown in shaded blue according to blastN legend. **c**, Growth curves of *E. coli* DH5α carrying EcCI^B25FEP^ or not (control) in M9 minimal medium with raffinose (30 mM, red lines), Glucose (30 mM, black lines) or without sugar as carbon source (grey lines). Source data are provided as Source Data file. **d**, EcCI^B25FEP^-encoded protection against phage T4 infection. Upper panel: Lysis plaque assays of 10-fold serial dilutions of T4 phage on *E. coli* DH5α carrying EcCI^B25FEP^ or not (control). Similar plaque assays of other phages are shown in Extended Data Fig. 8. Lower panel: Growth curves of EcCI^B25FEP^-carrying *E. coli* DH5α (solid) and control (empty) with or without infection by phage T4 at an MOI of 0, 0.05, or 5. Each curve represents the mean ± SD of five independent experiments. Source data are provided as Source Data file.

Moreover, elements of the EcCI^tRNA-Phe^ family were found to contain up to five different defense systems, the most prevalent being a type I restriction-modification system (TGANNNNNNCTTC; N6A), the recently described Ogmios system and the Mokosh type IA system (Extended Data Fig. 7, Supplementary Table 2)^20–22^. In addition to the RM and Ogmios systems, the prototype EcCI^B25FEP^ harbours other predicted anti-phage systems (i.e., PAPO_2678/2679, PAPO_5738/11762…) and three different Toxin-Antitoxin systems *higBA*, *cbeA*-*cbtA*, and *parDE* (Fig. 1a, Extended Data Fig. 7)^20,21^. While EcCI^B25FEP^ did not seem to inhibit infection by its helper jumbo phage Beqaa (Extended Data Fig. 6), we however screened the EcCI^B25FEP^-encoded anti-phage defense against different lytic *Escherichia* phages^23^. Chromosomally-integrated EcCI^B25FEP^ conferred a broad anti-phage protection against different lytic *Myoviridae* and *Siphoviridae*, exhibiting very strong resistance to phage T4 (∼10^5^-fold decrease in lysis plaque assays) (Extended Data Fig. 8). This strong resistance phenotype was confirmed in time-dependent *E. coli* infection by phage T4 in presence/absence of EcCI^B25FEP^ (Fig. 2d).

The PICJI family represents the larger phage satellites to date, with EcCI^B25FEP^ being the largest satellite for which an initial deciphering of its life style has been carried out. EcCI^B25FEP^ satellite appeared highly dependent on its helper jumbo phage for activation and probably packaging. Moreover, the production cycle of the helper jumbo phage Beqaa seemed not significantly affected by the presence of the active PICJI EcCI^B25FEP^. Such characteristics are shared with other satellites like the minimalist PICMIs which have also virulent helper phages^12^. In contrast and coherent with their larger size, PICJIs harbour a broad accessory gene repertoire dedicated to AMR and host adaptation, as well as anti-viral defense towards competitors of the helper phage^6^. Our data illustrate the highly complex interactions between satellites, helper phages, and bacterial hosts extending from parasitism to mutualistic beneficial^1,5^. Since a decade, there is a growing interest in nucleus-forming jumbo phages for phage therapy because they are virulent and resistant to most well-known defense systems^7,24^. Two concomitant studies recently published have revealed that several *Chimalliviridae* are virulent phages able to persist with their bacterial hosts^13,14^. In addition, our study highlights that the jumbo *Goslarvirus* Beqaa is also able to transduce efficiently the PICJI EcCI^B25FEP^ that spreads horizontally numerous cargo genes involved in anti-phage defense, host adaptation, and not least critically important AMR genes. Thus, their future use as therapeutics should be taken with caution^7^. This first study paves the way to understand the molecular complex interactions between helper jumbo phages and PICJIs (induction of excision, replication, packaging) during their common life cycle and more broadly to their evolutionary interplay and ecology.

## Methods

### Bacterial strains, growth conditions, and antimicrobial susceptibility testing

The *E. coli* field strains B25FEP & M24FEP have been described previously^8^. A full list of strains and plasmids used in this study is provided in Supplementary Table 1. Strains were routinely grown at 37°C in liquid or solid Luria Bertani (LB) medium, supplemented with appropriate selection as required: ceftriaxone (2 μg ml^-1^), kanamycin (50 μg ml^-1^), gentamicin (20 μg ml^-1^) and rifampicin (125 μg ml^-1^). Antibiotic susceptibility and the production of extended-spectrum β-lactamase were determined by the disc diffusion method on Mueller–Hinton agar and using the double-disc synergy test, respectively, as recommended by EUCAST (http://www.eucast.org/) and previously described^8^.

### Raffinose metabolism

The growth ability conferred by the *raf* operon of EcCI^B25FEP^ was assessed in M9 minimal medium (47.8 mM Na_2_HPO_4_, 22 mM KH_2_PO_4_, 8.6 mM NaCl, 18.7 mM NH_4_Cl, 2 mM MgSO_4_ and 100 μM CaCl_2_) supplemented with raffinose at 30 mM (Sigma-Aldrich), as the sole source of carbon. Overnight cultures of *E. coli* DH5α containing EcCI^B25FEP^ or not were washed twice in M9 minimal medium and diluted 1:100 in M9 supplemented with 100 nM thiamine and 0.02 % casamino acids, and either 30 mM glucose (positive control), 30 mM raffinose, or without sugar carbon source (negative control). Optical density at 600 nm (OD_600_) was monitored at 37°C every 20 min for 840 min in microplates (5 replicates) using the Bioscreen C MBR^TM^ system (Thermo Fisher). Growth curves were produced using ggplot2 in RStudio (v2026.01.1+403).

### Phage isolation and enrichment, and phage infection time-course assays

The jumbo phage Beqaa was concomitantly isolated with the expanded-spectrum-cephalosporin resistant *E. coli* strain B25FEP from a chicken rectal swab in 2018 in Lebanon^8^. Briefly, the initial isolation of the jumbo phage Beqaa was carried out from 200 ml LB overnight culture of *E. coli* strain B25FEP which was centrifuged twice at 3,500g for 20 min. The resulting supernatant containing phage particles was purified following the NaCl/PEG-8000 enrichment protocol below. To obtain pure lysate of vB_EcoM_Beqaa devoid of phage particles containing EcCI^B25FEP^, serial dilutions of the initial phage lysate were amplified on sensitive *E. coli* DH5α and purified by NaCl/PEG-8000 precipitation. The absence of EcCI^B25FEP^ containing phage particles was confirmed by PCR using specific primers (Supplementary table 3) as well as by the absence of EcCI^B25FEP^ transfer in transduction assays (see below).

#### Phage purification and concentration by NaCl/PEG-8000 precipitation

Firstly, supernatants of bacterial cultures were treated with DNase I and RNase A (1 μg ml^-1^) at 37°C for 30 min. After filtration through a 0.45 μm filter, 25 % (*w*/*v*) NaCl 2.5 M/PEG-8000 (Euromedex) was added to the phage lysate, briefly vortexed and precipitated at 4°C overnight. After centrifugation at 12,000g for 20 min at 4°C, the supernatant was discarded and the precipitate was resuspended in 4 ml TBS (Tris 20 mM, NaCl 150 mM, Sigma-Aldrich). A second precipitation was performed with 20 % NaCl 2.5 M/PEG-8000 for 1h on ice, then centrifuged at 16,000g for 10 min at 4°C. The supernatant was removed and the precipitate was resuspended in 600 μl TBS and filtrated at 0.22 μm before storage at 4°C or 20°C depending on subsequent applications.

#### Crude phage extract

Briefly, a LB agar plate was inoculated with 1:100 dilution of overnight liquid culture of a sensitive *E. coli* strain and then dried for 15-30 min at room temperature. Then, 100 μl of phage lysate was applied at the middle of the LB agar plate and dried for an additional 15 min. LB plate was incubated lid on top at 37°C overnight. The day after, the resulting lysis plaque was cut and placed in 20 ml LB broth and incubated at 37°C under shaking conditions at 200 r.p.m. for 4h. Then, 200 μl chloroform was added to the solution, then vortexed vigorously for 15 s, and incubated at room temperature for 1h. After centrifugation at 6,000g for 10 min, the supernatant was passed through a 0.45 μm filter and the resulting phage suspension was aliquoted and stored at 4°C.

For all phage lysates, the absence of bacteria was confirmed by incubating 10 μl of phage lysates in 5 ml LB broth at 37°C under shaking conditions at 200 r.p.m. for 48h before subsequent use.

Phage infection time-course assays were performed in liquid LB medium. Overnight cultures of the recipient *E. coli* strain DH5α containing EcCI^B25FEP^ or not (control) were diluted 1:500 in fresh LB medium and grown at 37°C and 200 r.p.m. to reach an OD_600_ of 0.20/0.22. Then, 200 μl of bacterial cultures (ca. 10^6^ bacteria) was infected with 100 μl of phage lysates at various MOI. OD_600_ was monitored at 37°C every 20 min for 5 technical replicates at least during 720 min using the Bioscreen C MBR^TM^ system (Thermo Fisher). Growth curves were produced using ggplot2 in RStudio (v2026.01.1+403).

### Plaque assays, EcCI^B25FEP^ transduction assays, host-range, and anti-phage protection of EcCI^B25FEP^

Two different methodologies were used in plaque assays to assess phage lysis efficiency. First, large square LB plates were inoculated with 3 ml of bacterial culture (OD_600_ ∼ 0.4) and the excess of bacterial culture was removed. After 30 min drying, plates were spotted with phage lysate dilutions. The double agar overlay plaque assay was also performed as previously described elsewhere^5^. Briefly, 520 μl of bacterial culture (OD_600_ ∼ 0.4) were added to 13 ml of top LB soft agar (0.5 %), then poured onto large square LB plate and let for 45 min drying. In both methodologies, plates were spotted with 4 μl serial 10-fold dilutions of phage lysates (10^0^ to 10^-8^). Plates were incubated at 37°C overnight and then imaged.

Bipartite transduction assays of the PICJI EcCI^B25FEP^ were carried out in a similar manner as conjugative mating experiments in liquid cultures using sodium azide-resistant (Az^R^) *E. coli* K-12 J5-3 as recipient strain and EcCI^B25FEP^-containing *E. coli* field strains in presence/absence of the jumbo phage vB_EcoM_Beqaa as donors, as previously described^9^. Transductants were selected on LB plates containing ceftriaxone at 2 μg ml^-1^ and sodium azide at 250 μg ml^-1^. Transductants were screened by PCR for the presence of EcCI^B25FEP^, using primers in Supplementary Table 3.

To precisely determine EcCI^B25FEP^ transduction efficiency from phage lysates, the recipient *E. coli* DH5α was grown in 2 ml LB until 0.6 OD_600_, and then 30 μl of NaCl/PEG-precipitated phage lysate was added and incubated at 37C° for 1h. Ten-fold serial dilutions of the phage-bacterial infection mixture were spread on LB plate containing ceftriaxone at 2 μg ml^-1^ and incubated overnight at 37°C to enumerate EcCI^B25FEP^ transductants. The host range of EcCI^B25FEP^ transduction by Beqaa was determined for bacterial strains listed in Supplementary Table 1 using B25FEP phage lysate and the above transduction assay.

The anti-phage defense conferred by EcCI^B25FEP^ was determined in *E. coli* DH5α, first by plaque assays using different virulent *Escherichia* phages^23^ including model phages T4 and T7 as well as by phage infection time-course assay for the phage T4 (MOI= 0.05 and 5) as described above.

### Transmission electron microscopy

Briefly, 10 μl of NaCl/PEG purified phage lysates or *E. coli* DH5α cultures infected with the jumbo phage Beqaa were deposited onto carbon film-covered copper grids. After 1-2 min, excess liquid was blotted off with filter paper. For staining, 3 drops of 1.5% (w/v) aqueous uranyl acetate solution were successively deposited then blotted off, after which the grid was allowed to dry under ambient air. The negatively stained samples were then imaged at 120 kV using a JEOL JEM-1400 Plus transmission electron microscope equipped with an AMETEK-Gatan OneView CMOS camera. Averages tail length and head diameter were determined from 100 phage particles using ImageJ v1.54g.

### Gene inactivation, cloning and quantitative PCR

Deletion of the integrase gene *intS* and the excisionase gene *alpA* was performed in *E. coli* DH5α containing EcCI^B25FEP^ by the one-step chromosomal gene inactivation technique of Datsenko & Wanner^25^. Briefly, the kanamycin resistance gene *kan* flanked by FRT sites was amplified by PCR using the template plasmid pKD4 and hybrid primers for recombination (Supplementary Table 3). These primers were synthesized with 20 nucleotides of priming sites of pKD4 and with 50 nucleotides upstream and downstream of the start and stop codon of the ORF, respectively. The 1.6 kb PCR product was purified and electroporated into *E. coli* DH5α_EcCI^B25FEP^ expressing the λ Red recombinase from plasmid pKD46-Gm^26^. Homologous recombination between the PCR product and the genomic DNA resulted in the deletion of the ORF and its replacement by the *kan* resistance cassette. PCRs were performed to confirm that the EcCI^B25FEP^ mutant strains had the deletion correctly located (Supplementary Table 3). To eliminate the kan cassette, the EcCI^B25FEP^ mutant strains were transformed with the FLP recombinase expression plasmid pCP20-Gm^26^, and gentamicin-resistant transformants were selected at 30 °C. After culture at 37 °C and/or 43 °C, a large majority of transformants had lost the FRT flanked kan gene and also the FLP helper plasmid pCP20-Gm and thus became susceptible to both kanamycin and gentamicin. The resulting mutant strains named DH5α_EcCI^B25FEPΔ^*^intS^* and DH5α_EcCI^B25FEPΔ^*^alpA^* were confirmed by PCR using primers flanking the deleted region and nucleotide sequencing revealed that it harboured a 85 nucleotides scar in place of the ORF as expected.

Plasmidic vector pBAD18-kan was used to directionally clone the *intS* and *alpA* ORFs of EcCI^B25FEP^ under the control of the arabinose *P_BAD_* promoter (Supplementary Table 1). Briefly, ORFs were amplified by PCR using the Phusion High-Fidelity DNA polymerase (New Englands Biolabs) and specific primers (Supplementary Table 3). The PCR product was cloned into the pCR2.1 TA cloning vector (Invitrogen®). By *Eco*RI-*Xba*I digestion, the inserts of the recombinant plasmids carrying the *intS* and *alpA* ORFs were further subcloned into pBAD18-kan, resulting in plasmids named pintS and palpA, respectively.

Excision rate as well as copy number of EcCI^B25FEP^ (circular and integrated) were assessed by real-time quantitative qPCR. Primers used for quantification are listed in Supplementary Table 3. The data were analyzed using the threshold cycle (ΔΔ*C_T_*) method using the 3 chromosomal target genes *dnaB*, *hicB* and *dsbD* as chromosomal references. Total copy numbers of EcCI^B25FEP^ and vB_EcoM_Beqaa were assessed using *rafA* and *phuZ* as target genes, respectively. Excised and integrated forms of EcCI^B25FEP^ were assessed using the *attP* site of the excised form and the left junction with *tRNA-Phe* as the targets, respectively.

### DNA extraction and whole-genome sequencing

Genomic DNA of bacterial strains were extracted from 1 ml liquid culture using the NucleoSpin tissue kit following the manufacturer’s recommendations (Macherey-Nagel, Düren, Germany). Phage lysates were previously treated with DNase I (1 μg ml^-1^) at 37°C for 30 min to remove free bacterial DNA before phage DNA extraction using the Monarch® HMW DNA extraction kit (*BioLabs*_Inc_). DNA samples were sent to Genewiz^®^ From Azenta Life Sciences (Leipzig, Germany) for whole-genome sequencing on Illumina NovaSeq 6000 system using 2 x 150-bp paired-end sequencing. Raw reads were quality filtered, and sequencing adapters were removed by Genewiz^®^ before data delivery. Long-read sequencing was realized in house using Oxford Nanopore Technlogies (ONT, Oxford, U.K.). Briefly, bacterial or phage DNAs were barcoded and multiplexed using the native-barcoding kits following the manufacturer’s recommendations. Long-read sequencing was performed using the MinION sequencing device Mk1B and flowcell R9.4.1 or R10.4.1. Flowcells, kits, sequencing runs, and basecalling characteristics are indicated in Supplementary Table 4.

### Sequence analysis

Complete genomes of bacterial strains as well as phages were obtained through hybrid assembly using the FullForce Plasmid assembler (https://github.com/NorwegianVeterinaryInstitute/FullForcePlasmidAssembler). Briefly, Oxford Nanopore raw reads were quality-filtered at >Q10 and trimmed using QCAT. Illumina raw reads were trimmed using trimmomatic v.032 and hybrid assemblies were realized using Unicycler v0.4.7.

Whole-genome comparisons were firstly carried out by multiple genome alignments using Mauve to localize horizontally transferred DNA between donor strains and transductants. The GenoFig visualization tool v1.1.0 and blastN algorithm were used to identify precisely chromosomal integration sites, to create genetic maps of EcCI^B25FEP^ and the jumbo phage vB_EcoM_Beqaa, and to compare various EcCI^B25FEP^-related elements^27^. EcCI^B25FEP^-related elements were searched in prokaryotic complete genomes of the GenBank database using the 72,178-bp query sequence of *tRNA-Phe*::EcCI^B25FEP^ on February, 18^th^ 2026. BlastN results were cleaned to keep only subject sequences corresponding to EcCI^B25FEP^-related elements integrated at *tRNA-Phe* and extending from direct repeat sequences *attL* and *attR*, resulting in 273 sequences that were retrieved from NCBI GenBank database. To establish evolutionary relationships between these EcCI^B25FEP^-related elements, the pangenome analysis tool PPanGGOLiN v2.2.6 was used to determine softcore genes shared among the EcCI^B25FEP^ family based on the 273 sequences as identified above and the EcCI^B25FEP^ sequence added as reference^28^. Then, softcore genes were concatenated and then aligned using mafft v7.526, and the resulting alignment was provided to IQ-TREE v3.0.1 to build a maximum likelihood phylogenetic tree using ModelFinder for model selection^29–31^. The phylogenetic tree was visualized and annotated with the iTOL web interface tool v7.5^32^. Determination of Sequence-Type (ST), *Escherichia* phylogroups, *Salmonella* serotype, search for antimicrobial resistance genes, virulence-associated genes and defense systems were done using mlst v2.33.1, ClermonTyping v.23.6, SeqSero2 v1.3.2, ResFinder v4.7.2 and VirulenceFinder v3.2.0, DefenseFinder v3.0.0, respectively^20,21,33–37^.

Oxford Nanopore and Illumina raw reads resulting from the sequencing of the phage lysate DNA were mapped against the assembled genomes of the jumbo phage Beqaa, the EcCI^B25FEP^, and the recipient *E. coli* DH5α chromosome using minimap2 v2.28 and Bowtie2 v2.5.3, respectively^38,39^. Information of mapped reads (number/%, aligned bases, N50) were extracted to determine mean coverage and RPKM (reads per kilobase per million mapped reads). Mean coverages were plotted as a function of read N50 (kb) and RPKM for Oxford Nanopore and Illumina mapping, respectively, using ggplot2 in RStudio (v2026.01.1+403).

## Data availability

The complete sequence of nucleus-forming jumbo *Escherichia* phage vB_EcoM_Beqaa has been deposited in NCBI database under GenBank accession number PZ012685. All other sequencing data including raw reads and complete assembled genomes have been deposited in NCBI under Bioproject number PRJNA1424553. Any further information, strains and materials are available from the corresponding author upon reasonable request. Source data are provided with this paper.

## Acknowledgements

We dedicate this work to the memory of Professor Dolla Karam Sarkis (Deceased in 2023) who was involved in the initial phase of this study. She had been an exceptional mentor for microbiology students at the Saint Joseph University of Beirut in Lebanon. We also thank the electron microscopy facility for phage imaging (US61 Analyse des Systèmes Biologiques, Tours, France). In addition, we warmly acknowledge Dr. Catherine Schouler who provided *Escherichia* lytic phages as well as for thoughtful discussions. We also thank Thierry Cochard for his technical assistance in qPCR experiments.

## Author contributions

D.K.S. and B.D. designed, conceived and supervised the project. M.M. initially performed field strain isolation and first horizontal transfer experiments. A.R., M.R.A., K.P., I.F., and L.F. performed experimental molecular characterizations of the study. P.I.R. realized electron microscopy imaging. S.O.L., F.X.W. and B.D. performed genomic analyses. A.R., M.R.A. and B.D. wrote the original draft and editing of the manuscript. All other authors except D.K.S. provided feedback, review the manuscript and approved its final version.

## Funding

This study was funded by public funds from the National Research Institute for Agriculture, Food and Environment (INRAE) in France and co-funded by the European Union’s Horizon 2020 project 101136346, EUPAHW. Views and opinions expressed are, however, those of the author(s) only and do not necessarily reflect those of the European Union or the European Research Executive Agency. Neither the European Union nor the granting authority can be held responsible for them. M.M. was funded by an USJ PhD studentship (grant numbers FP72/73), by the National Center for Scientific Research in Lebanon (CNRS-L) and by the Francophone University Agency (AUF) in Lebanon during her stay in France.

---

**Extended Data Fig. 1.**
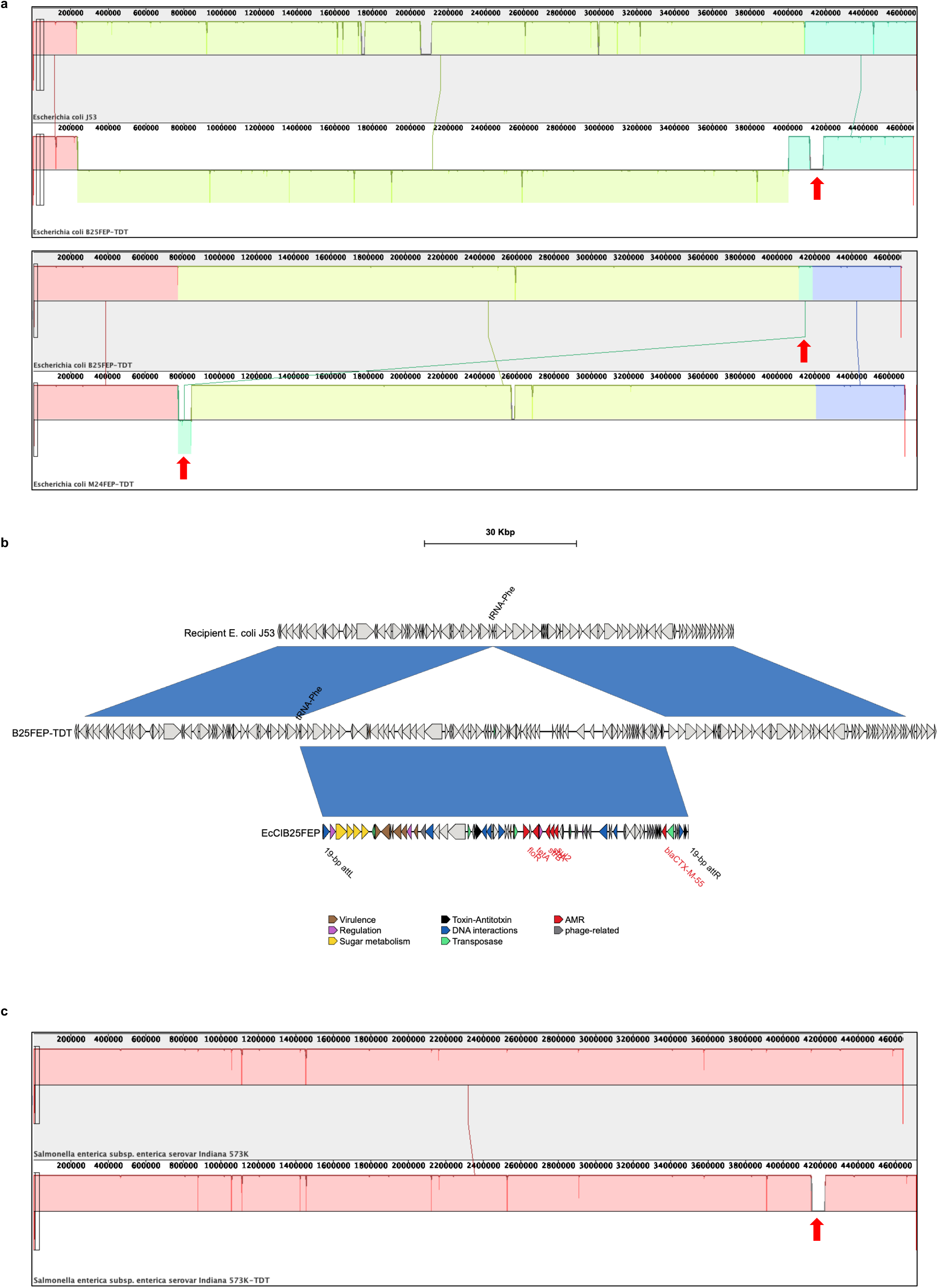
Chromosomal insertion sites of *bla*_CTX-M-55_-carrying islands in *E. coli* and *S.* Indiana recipient strains. **a,** Mauve comparisons of the chromosomes of the Az^R^ *E. coli* recipient strain J53 and transductants B25FEP-TDT and M24FEP-TDT. The red arrow shows the chromosomal integrated *bla*_CTX-M-55_-carrying islands. **b,** Zoom in the *tRNA-Phe* insertion site of EcCI^B25FEP^. Sequence homologies are shown in dark blue (100%) between adjacent sequences. **c,** Mauve comparisons of the chromosomes of the *S.* Indiana recipient strain 573K and its transductant carrying EcCI^B25FEP^ (red arrow).

**Extended Data Fig. 2.**
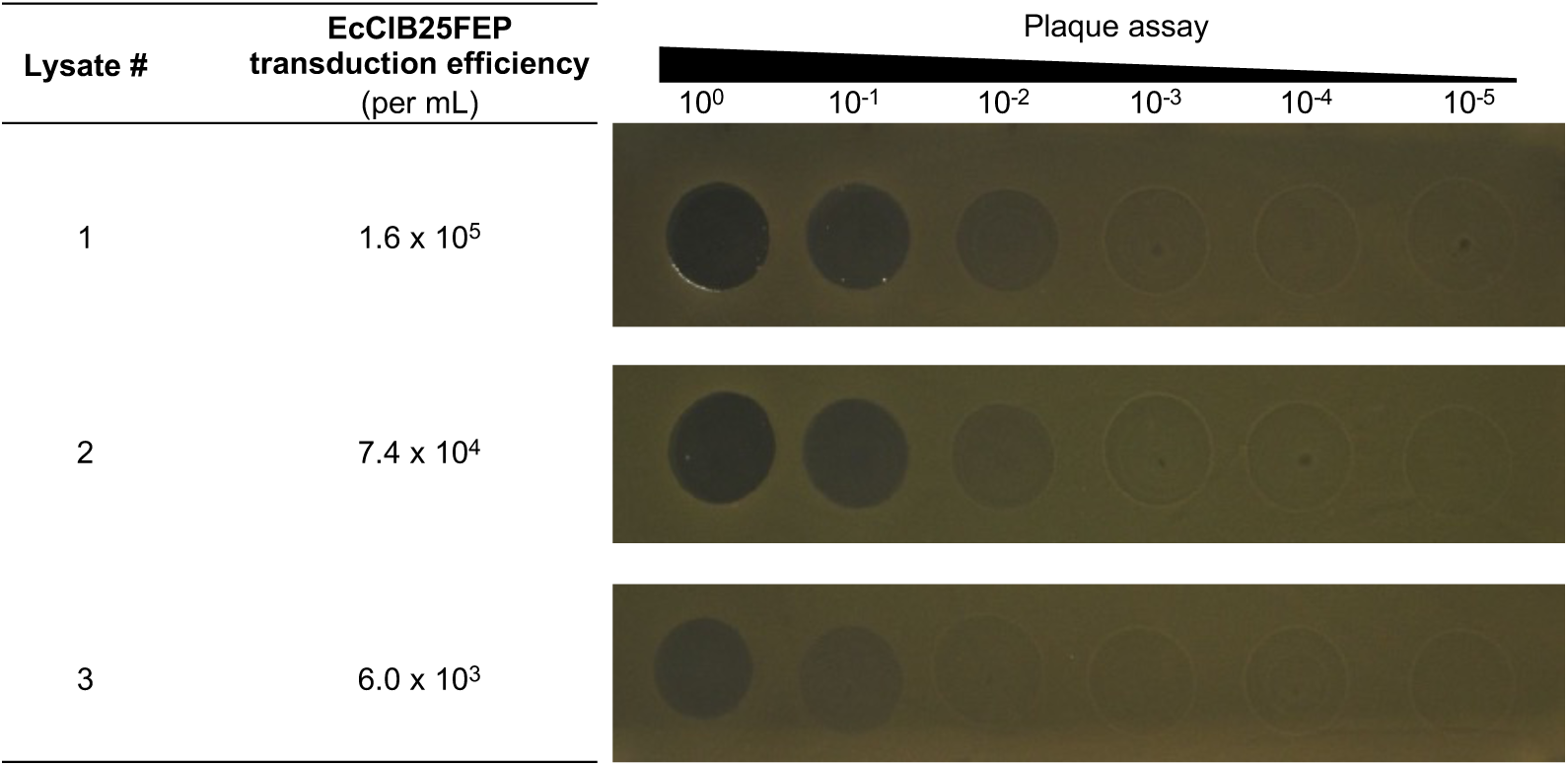
Correlation between EcCI^B25FEP^ transduction efficiency and *E. coli* DH5α infection efficiency. The transduction efficiency of EcCI^B25FEP^ into the recipient *E. coli* strain DH5α was assessed for 5 different phage lysates produced from the stock culture of the animal strain B25FEP (left panel). Concomitantly, lysis efficiency was determined by plaque assays on *E. coli* strain DH5α (right panel). Raw uncropped images are provided as Source Data files.

**Extended Data Fig. 3.**
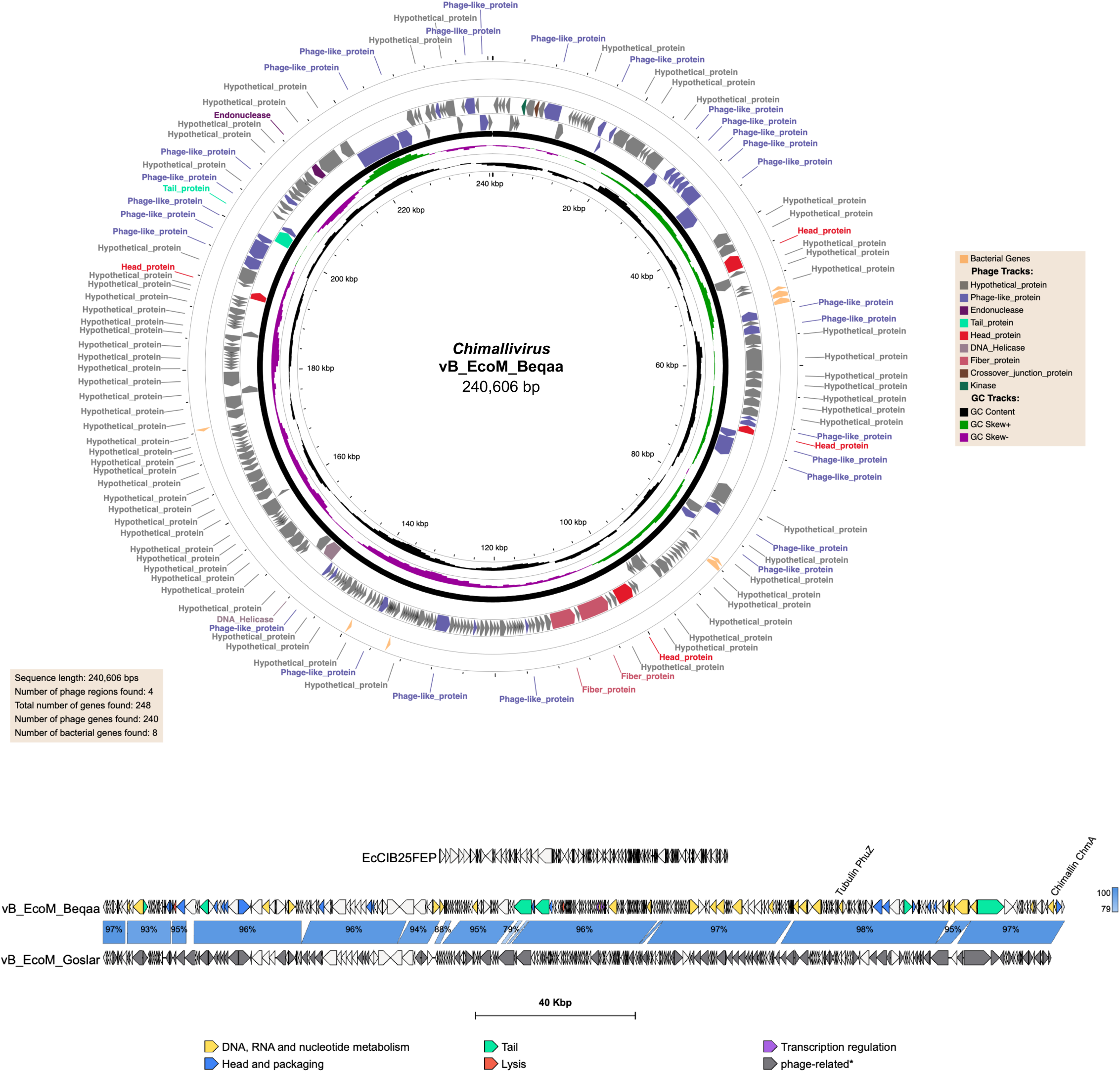
Genetic map and comparison of the Goslarvirus vB_EcoM_Beqaa. Upper panel, Sequence analysis, ORF prediction and genetic circular map were performed using the PHASTEST tool^40^. From inside to outside circles: genetic scale, GC content, GC skew, phage-related genes, and bacterial genes. ORFs are color-coded according to encoded-functions in legend. Lower panel, Genome comparison of the helper jumbo *Escherichia* phage vB_EcoM_Beqaa with the nucleus-forming *Goslarvirus* Goslar (NC_048170.1) of the *Chimalliviridae* family. EcCI^B25FEP^ is included in the figure to indicate the absence of homology with its helper phage. Sequence homologies larger than 500 bp are shown in shaded blue according to blastN legend. Both phages and EcCI^B25FEP^ were annotated with pharokka v1.9.1. Open reading frames of vB_EcoM_Beqaa are color-coded according to protein functions in legend.

**Extended Data Fig. 4.**
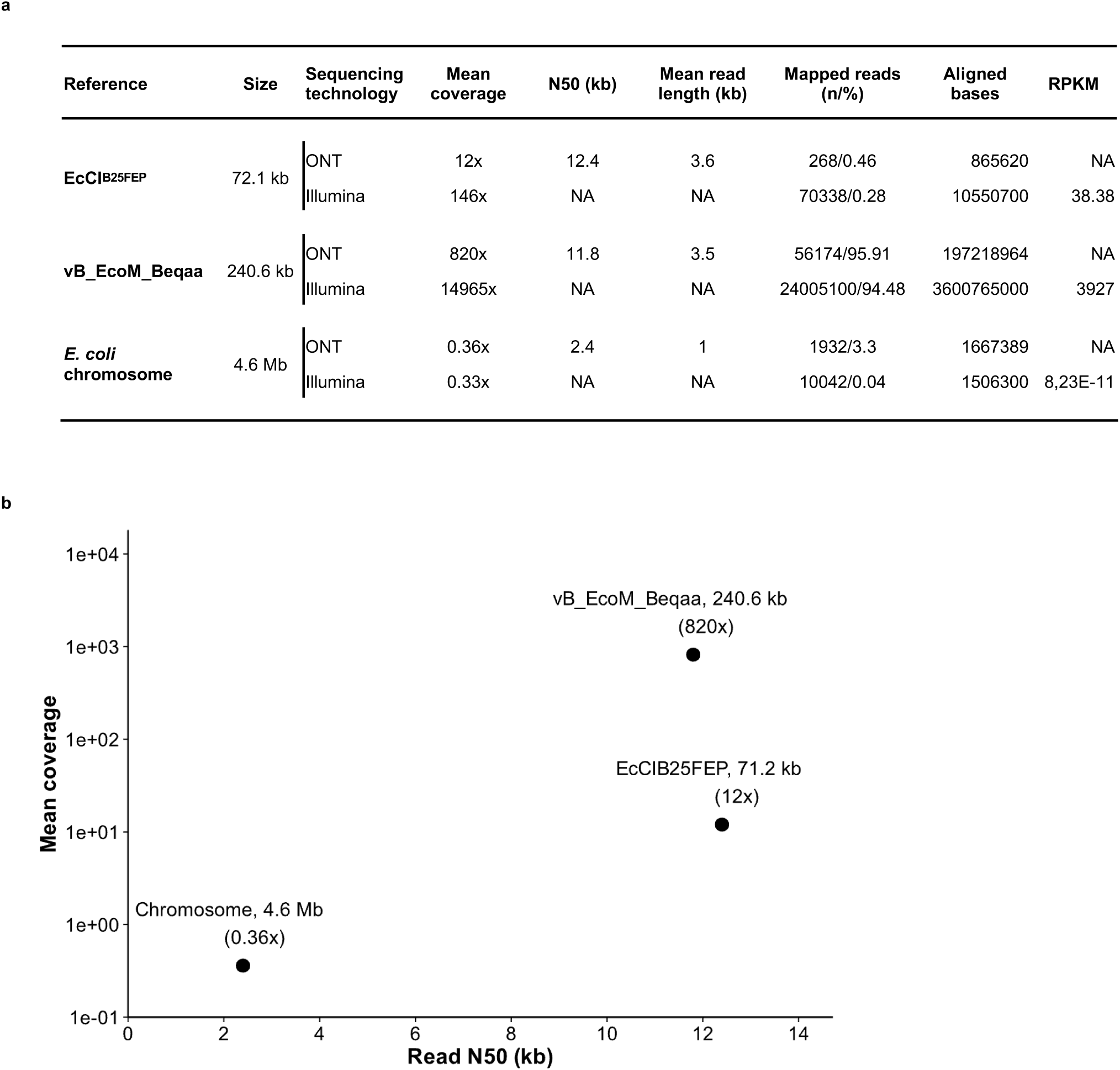
DNA sequencing of phage lysate DNA transducing EcCI^B25FEP^. **a,** Main mapping characteristics of Oxford Nanopore (ONT) and Illumina raw reads of phage lysate DNA. **b,** Scatter plot showing the relationship between N50 read length and mean coverage in Oxford Nanopore sequencing of phage lysate DNA for the phage vB_EcoM_Beqaa, the chromosomal island EcCI^B25FEP^, and the *E. coli* chromosome. Genome size and mean coverage are indicated for each point.

**Extended Data Fig. 5.**
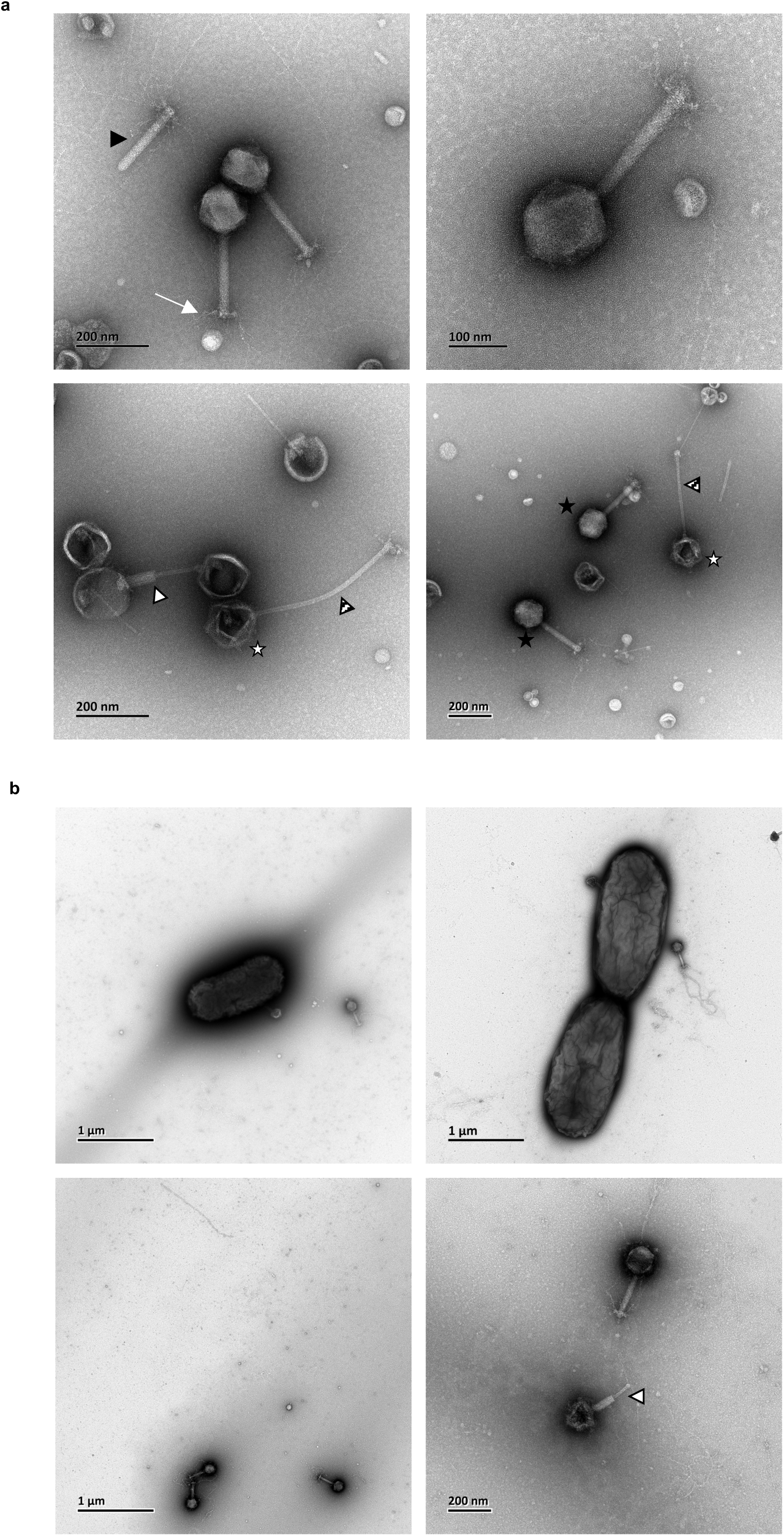
Transmission electron micrographs of the phage vB_EcoM_Beqaa. **a,** Phage particles from NaCl/PEG-purified lysates. **b,** *E. coli* DH5α culture infected by vB_EcoM_Beqaa. Samples were negatively stained with 1% uranyl acetate. Scale bars are included in each picture. Black, white and hatched arrowheads show broken, contracted tail sheath, and tail tube without sheath, respectively. White arrow highlights tail fibres. White and black stars show empty and full head capsids, respectively. Image files are provided as Source Data file.

**Extended Data Fig. 6.**
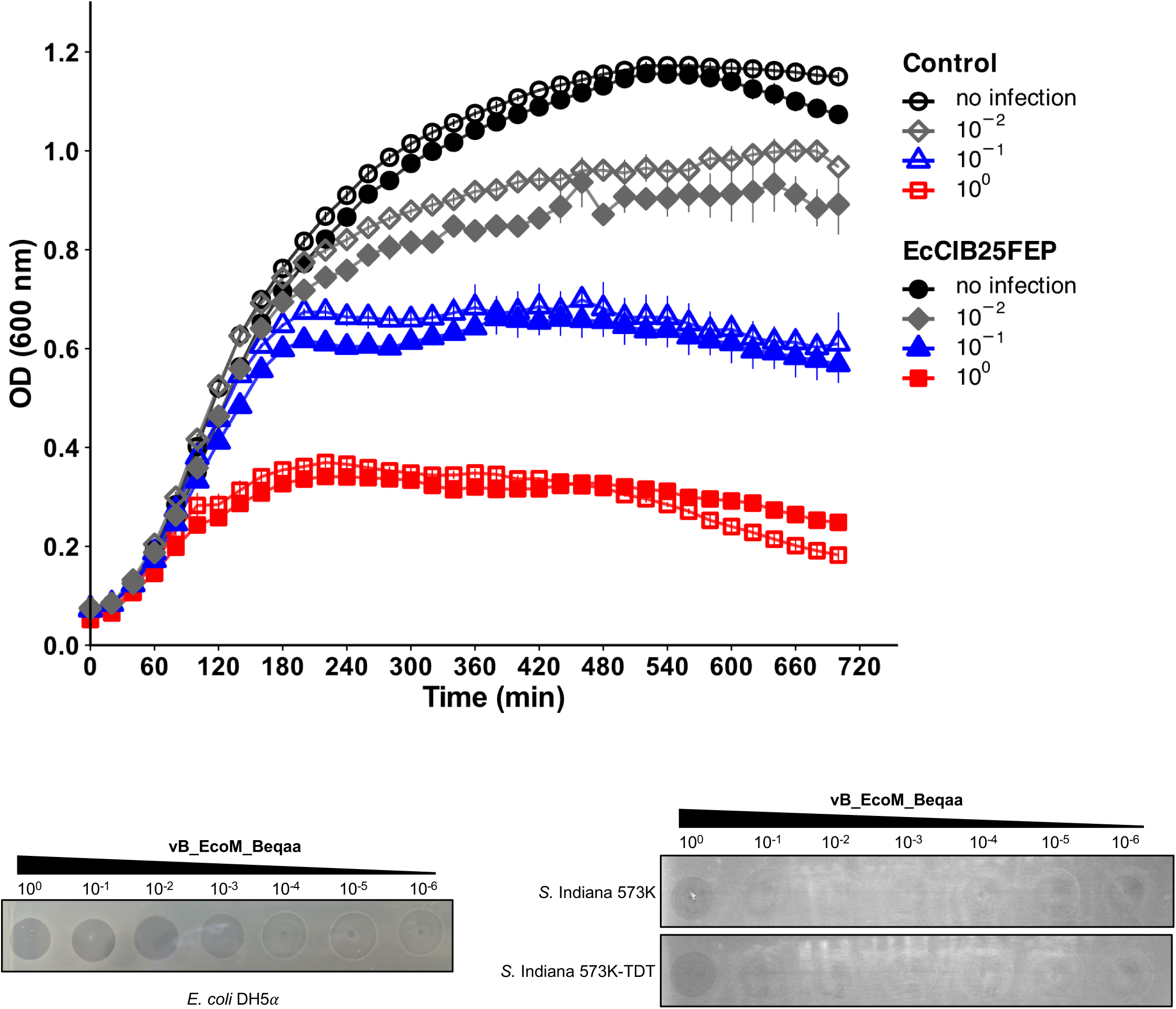
Infection of *E. coli* and *S.* Indiana carrying or not EcCI^B25FEP^ by vB_EcoM_Beqaa. Upper panel: Growth curves of *E. coli* strain DH5α carrying EcCI^B25FEP^ (solid symbols) or not (control: empty symbols) with or without infection by vB_EcoM_Beqaa. Infections were performed with 10-fold serial dilutions of phage vB_EcoM_Beqaa (See also Fig. 1,d). Each curve represents the mean ± SD of five independent experiments. Lower panel: Plaque assays of 10-fold serial dilutions of phage Beqaa lysate on *E. coli* strain DH5α (lower left, *S.* Indiana strain 573K and its transductant carrying EcCI^B25FEP^ (573K-TDT) (lower right). Raw uncropped image and source data are provided as Source Data files.

**Extended Data Fig. 7.**
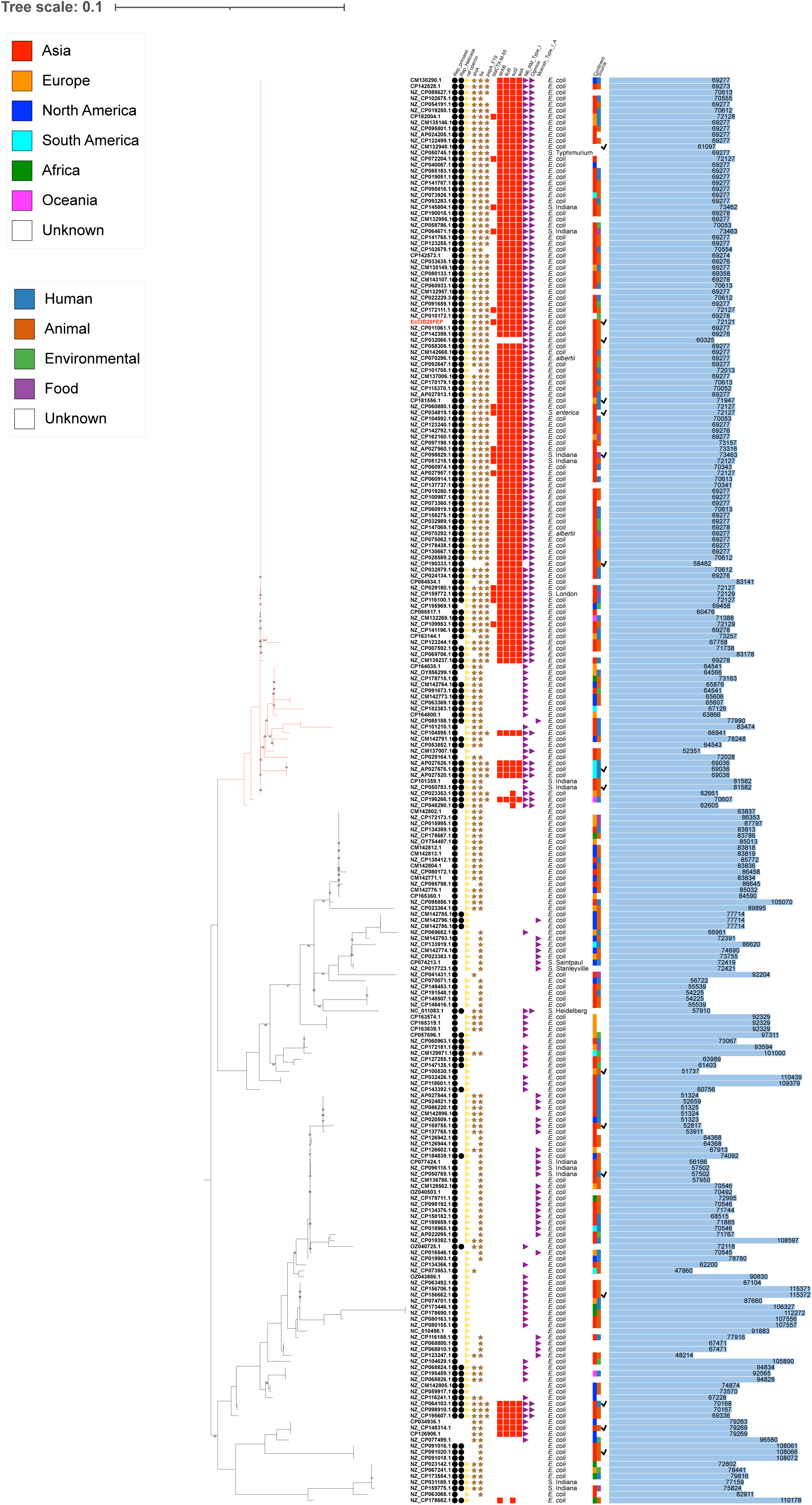
Phylogenetic relationship of 237 EcCI^B25FEP^-related chromosomal islands integrated at *tRNA-Phe*. (Supplementary Table 2). The tree is based on the alignment of soft-core genes (see methods) and rooted at mid-point. Branches of the largest cluster containing EcCI^B25FEP^ are coloured in red. In addition to GenBank accession numbers, the main EcCI^B25FEP^-encoded functions (replication, raffinose metabolism, virulence-associated factors, antibiotic resistance, defense systems), bacterial host species, geographical origin, source, and size, are shown on the right of the tree. Genetic organisation of EcCI^B25FEP^-related elements marked with a tick are fully compared in Fig. 2,b.

**Extended Data Fig. 8.**
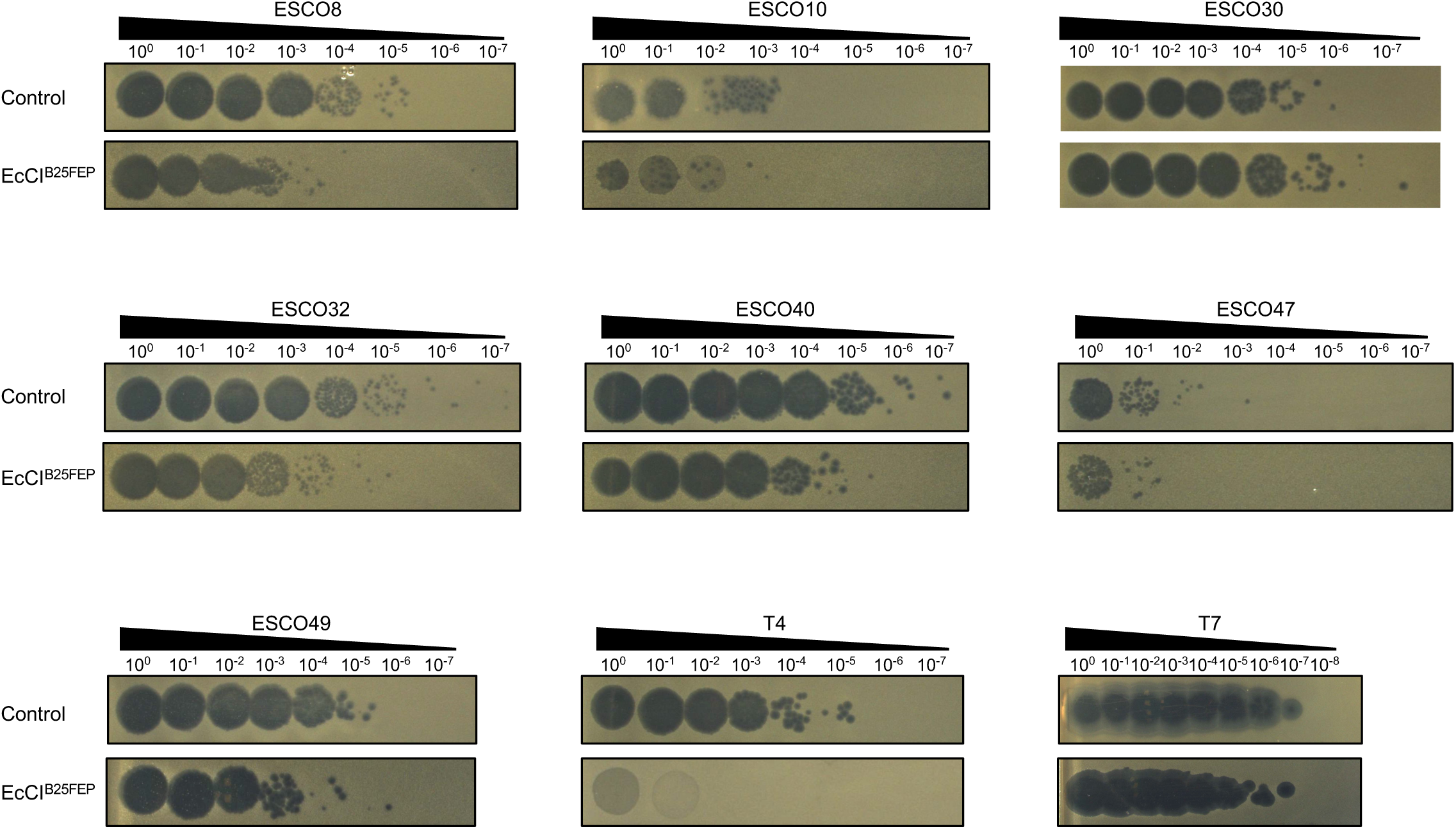
Lysis plaque assays of 10-fold serial dilutions of different lytic *Escherichia* phages on *E. coli* strain DH5α carrying EcCI^B25FEP^ or not (control). Raw uncropped images are provided as Source Data files.

## Notes

### Competing Interest Statement

The authors have declared no competing interest.

